# Dysregulated lactate metabolism synergizes with ALS genetic risk factors to accelerate motor decline

**DOI:** 10.1101/2025.11.24.690227

**Authors:** Shweta Tendulkar, Tong Wu, Amy Strickland, Amber R. Hackett, Yurie Sato-Yamada, Xianrong Mao, Yo Sasaki, Jeffrey Milbrandt, A. Joseph Bloom, Aaron DiAntonio

**Affiliations:** Department of Developmental Biology, Washington University School of Medicine, St. Louis, 63110, United States; Department of Genetics, Washington University School of Medicine, St. Louis, 63110, United States; Center for Advanced Oral Science, Niigata University, Niigata Japan; Needleman Center for Neurometabolism and Axonal Therapeutics

## Abstract

Neurons rely on glial ‘lactate shuttling’ for metabolic support, which declines with aging and in neurodegenerative disease. Full disruption of lactate shuttling in peripheral nerves causes progressive axon degeneration, but we were interested to understand how partial disruption, a scenario more relevant to aging and disease, contributes to neurodegeneration risk. Pyruvate and lactate are interconverted by lactate dehydrogenases (LDHA and LDHB) in both lactate producing and consuming cells. We therefore began by investigating *Ldhb* knockout mice (loss of LDHA, the dominant LDH in liver and muscle, caused embryonic lethality), and discovered that they develop progressive neuromuscular junction atrophy and functional decline without axon degeneration. Because even *Ldhb^+/-^* heterozygosity significantly affects motor behavior, we also wondered about a potential link to congenital disease and pursued this by identifying rare loss-of-function LDHB variants among ALS patients. Next, to better understand how LDHB loss leads to motor decline, we selectively deleted it in defined cell types. SC-specific deletion caused robust motor defects, whereas motor neuron–specific deletion has little effect. Reasoning that neuronal LDHB deficiency could model age-associated decline in lactate metabolism, we asked whether it would interact with ALS genetic risk. Indeed, motor-neuron LDHB deficiency synergizes with relatively mild ALS risk variants— *TDP43^Q331K^*and *Sod1^D83G^* knock-in alleles—to produce early motor neuropathy, indicating that LDHB loss enhances disease risk. These findings establish lactate metabolism as a modifier of motor system vulnerability and highlight it as a therapeutic target in peripheral as well as central neurodegeneration.

## Introduction

Neurons depend on tightly regulated metabolic support from surrounding glial cells, and disruption of this partnership is associated with profound health consequences. A main pillar of this support system is the ‘lactate shuttle’ by which glycolytic glia supply lactate to neurons, which then convert lactate back into pyruvate to drive the TCA cycle to support their high metabolic needs. The disruption or decline of lactate shuttling from astrocytes to neurons in the central nervous system (CNS)^1–3^ is implicated in both normal brain aging and in disease, including Alzheimer’s disease and ALS^4–6^. For example, astrocyte-motor neuron lactate shuttling is impaired and spinal cord lactate levels are reduced in both ALS mouse models and in ALS patients^7^. In both lactate producing cells and in consuming cells, lactate/pyruvate metabolism depends on Lactate Dehydrogenase (LDH), a multimeric enzyme consisting of two largely interchangeable subunits, LDHA and LDHB. These differ significantly in their expression patterns and are reported to differ in their substrate affinities, but both subunits robustly catalyze pyruvate/lactate interconversion in both directions. Blocking the lactate shuttle in peripheral nerves by deleting both LDH subunits causes progressive motor axon degeneration^8^ and there is little difference between knocking out LDH in motor neurons (MNs) themselves or in the Schwann cells (SCs) that surround and support motor axons.

Our earlier findings demonstrated that lactate shuttling is necessary for peripheral motor axon maintenance. However, complete ablation of the shuttling mechanism poorly represents the chronic decline that coincides with normal aging. We hypothesized that modest disruptions of lactate metabolism would also produce measurable phenotypes and exacerbate degeneration when neurons are otherwise stressed. In pursuit of a better model of lactate metabolism deficiency, we first examined whole-body LDHB knockout mice and found that, in contrast to the motor axon degeneration that results from completely abolishing SC-axon lactate shuttling, peripheral nerve axons are preserved when only LDHB is deleted. That is, endogenous expression of LDHA in SCs or motor axons is sufficient to prevent the spontaneous degeneration of motor axons. Nevertheless, whole-body LDHB knockout does result in progressive degeneration of neuromuscular junctions (NMJ). To clarify which cell types contribute to this defect, we therefore selectively deleted LDHB in neurons or Schwann cells and were surprised to discover that motor neuron LDHB knockout causes little harm, showing only very mild NMJ denervation in older animals. We reasoned that this modest disruption to lactate metabolism could serve as a genetic model of the reduction in glial-axon lactate shuttling that occurs with aging and in neurodegenerative disease. To test if mild lactate metabolism disruption can contribute to peripheral neurodegeneration risk, we combined mice with LDHB motor neuron KO with slowly progressing genetic models of ALS. We find that LDHB motor neuron loss synergizes with these pathogenic mutations in *TARDBP* and *SOD1*—to produce significantly enhanced and progressive motor impairment. In addition, we identify rare missense mutations in the *LDHB* gene in ALS patients and demonstrate that some encode dysfunctional LDHB protein. We conclude that declining lactate metabolism and shuttling may contribute to risk for peripheral neurodegeneration and suggest that targeting lactate shuttling is a candidate treatment strategy for neurodegenerative diseases.

## Results

### LDHB knockout mice have neuromuscular junction defects without axon loss

We originally developed murine *Ldha* and *Ldhb* genetic loss-of-function models to test the necessity of lactate shuttling in peripheral nerves^8^. Because LDHA and LDHB subunits both catalyze lactate/pyruvate interconversion in both directions, abolishing the lactate shuttling mechanism requires deleting both subunits from either the lactate-producing or lactate-consuming cells. In our prior study, we deleted both A and B in either Schwann cells, motor neurons, or sensory neurons using the Cre-Lox system^8^. With a goal of further dissecting the role of each subunit, we generated separate whole-body *Ldhb knockout* and *Ldha knockout* animals. *Ldha* expression dominates in liver and muscle tissues (**Fig. 1A**), and loss of *Ldha* causes embryonic lethality.^9^ However, *Ldhb* knockout mice appear grossly normal, healthy and fertile until at least 1 year of age.

**Figure 1:**
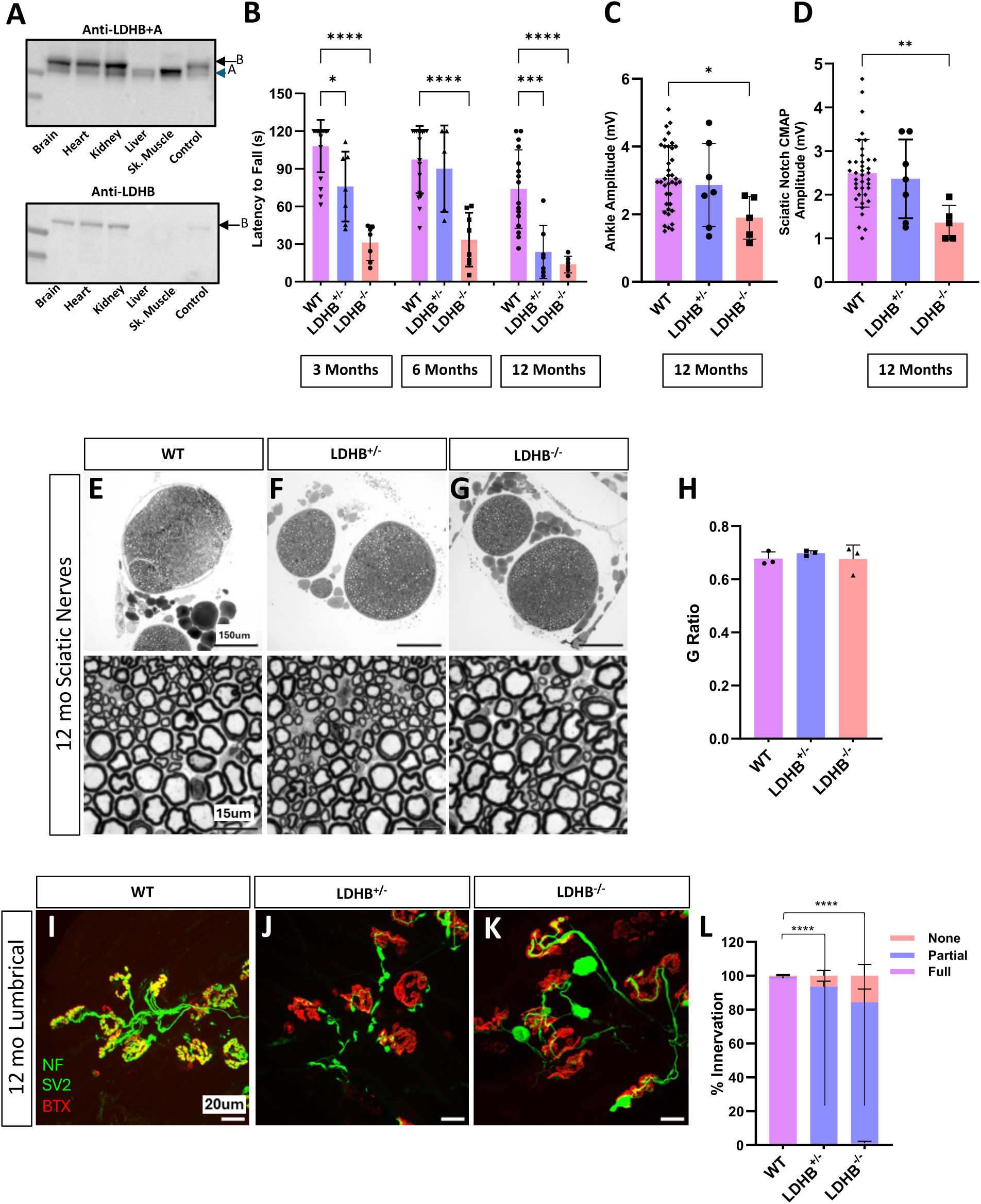
Whole-body knockout of LDHB causes progressive motor defects and NMJ defects but does not cause axon degeneration. Western blot analysis of LDH expression in multiple mouse tissues using antibodies that recognize both LDHA and LDHB (upper blot) or only LDHB (lower) **(A).** Motor function was measured in 3, 6 and 12-month-old wildtype (WT), *Ldhb^+/-^,* and *Ldhb^-/-^* animals using latency to fall from an inverted screen (n= 5-30 per genotype) **(B).** Compound muscle action potential (CMAP) measured at the ankle and sciatic notch in 12-month-old animals (n= 5-30) **(C** & **D).** Representative 10x and 100x images of Toluidine Blue-stained images of 12-month-old sciatic nerves from WT, *Ldhb^+/-^,* and *Ldhb^-/-^* animals **(E-G)** and quantification of the g-ratio for each (n=3) **(H).** Representative neuromuscular junctions on lumbrical muscles from 12-month-old WT, LDHB^+/-^, and LDHB^-/-^ animals stained in green to detect the synaptic vesicle marker synaptic vesicle glycoprotein 2A (SV2) and axon marker neurofilament medium chain (NF) and in red to detect post-synaptic endplates with bungarotoxin (BTX) **(I-K).** Percentage of innervation determined by colocalization of presynaptic SV2 with postsynaptic BTX divided into three categories: none (no overlap), partial, and full (n= 3-5; statistics are shown for comparisons of percent of fully innervated endplates) **(L).** ****p<10^-4^, ***p<0.001, **p<0.01, *p<0.05.

By 1 year of age, *Ldhb* knockout animals develop significant progressive motor behavior dysfunction and reduced compound muscle action potential (CMAP) amplitude (**Fig. 1B-D**) reminiscent of what we previously observed with either Schwann cell or motor neuron-specific knockout of both LDH subunits. However, surprisingly, these defects were not accompanied by axon degeneration (**Fig. 1E-H**). Instead, we observed significant neuromuscular junction (NMJ) defects without axon loss (**Fig. 1I-L**). Mild NMJ defects were also observed in whole-body *Ldhb^+/-^* heterozygous mice, although CMAP appeared normal, indicating a dose dependent effect. Thus, LDHB is not required to maintain axons, but proper NMJ maintenance requires full LDHB expression, most likely in the motor neurons or in Schwann cells.

Thus, to determine which cell types are responsible for the motor dysfunction observed in LDHB knockout mice, we used the Cre-Lox system to delete *Ldhb* only in motor neurons or only in Schwann cells. LDHB loss from MNs (LDHB MNKO) alone did not cause motor defects, but Schwann cell LDHB knockout mice (LDHB SCKO) developed motor phenotypes very similar to whole-body knockouts (**Fig. 2A-B**) implicating Schwann cells as the primary driver of the phenotype. LDHB MNKO mice do, however, exhibit mild denervation of lumbrical muscles by 1 year of age (**Fig. 2C-D**), suggesting long-term NMJ maintenance requires LDHB expression in both Schwann cells and motor neurons.

**Figure 2:**
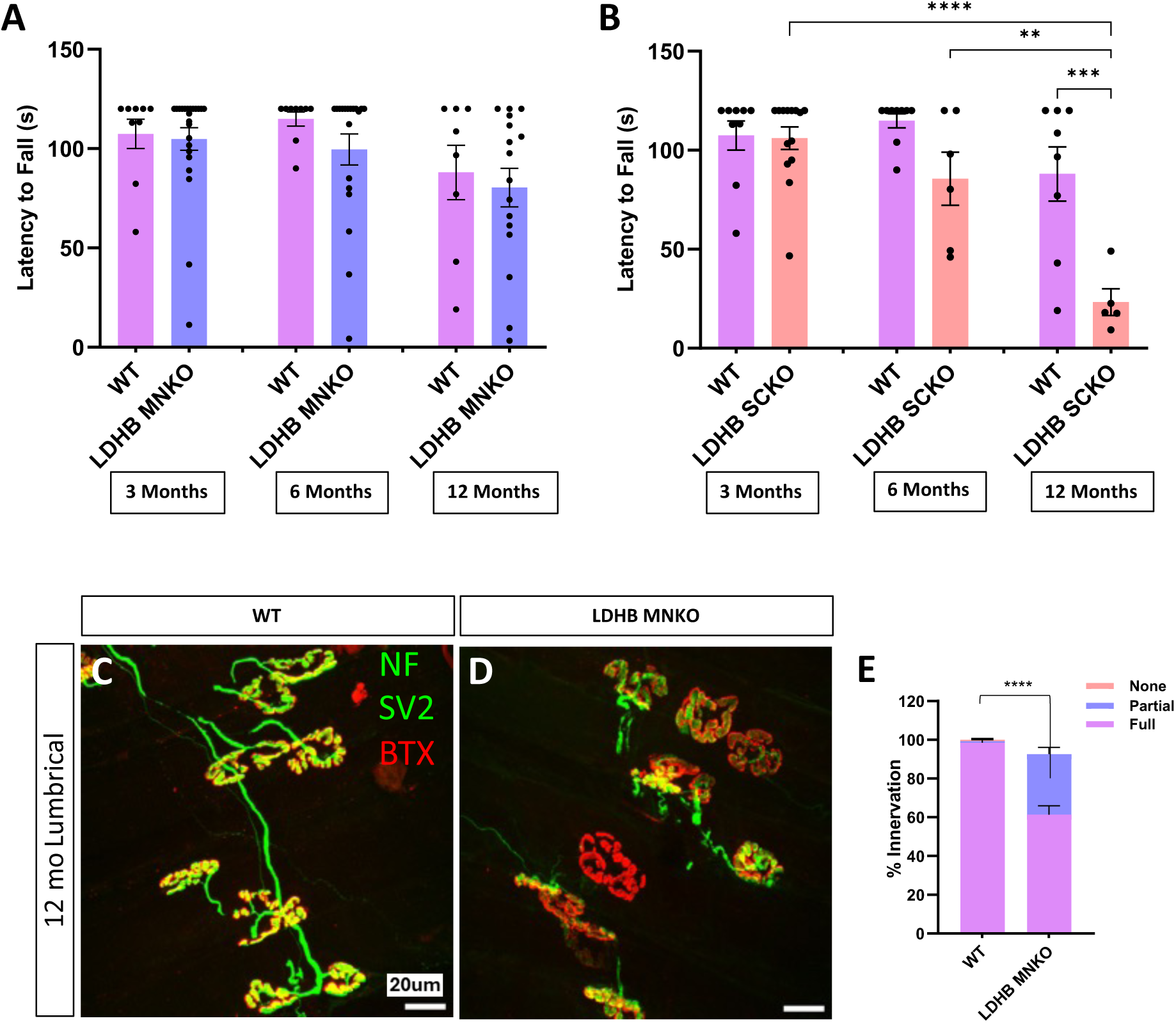
Knockout of LDHB in Schwann cells, but not in motor neurons, produces motor defects by 6 months old. Motor function (latency to fall from an inverted screen) measured in 3, 6 and 12-month-old WT, LDHB MNKO and LDHB SCKO animals (n= 15-20) **(A** & **B).** Representative images of neuromuscular junctions on lumbrical muscles from 12-month-old WT and LDHB MNKO animals stained with NF, SV2 and BTX **(C** & **D).** Percentage of innervation determined by colocalization of presynaptic SV2 with postsynaptic BTX divided into three categories, none (no overlap), partial, and full (n= 3; statistics shown for comparisons of fully innervated endplates) **(E).** ****p<10^-4^, ***p<0.001, **p<0.01.

### Rare *LDHB* loss-of-function alleles occur in ALS patients

Because whole-body *Ldhb^+/-^* heterozygous mice also develop progressive motor behavior and NMJ defects (**Fig. 1**) while appearing otherwise healthy, we wondered whether *LDHB* haploinsufficency might be an unappreciated risk factor for progressive peripheral neuropathy in human patients. We previously successfully pursued a similar question about rare variation in the *SARM1* gene and risk for peripheral neurodegeneration by combining genomic data from ALS case/control studies and enzymatic assays to determine the functional consequences of rare missense variants^10^. Therefore, to explore the disease risk associated with LDHB loss-of-function, we searched for rare (minor allele frequency <0.1%) *LDHB* missense and nonsense alleles in exome sequence data from several well-annotated studies of ALS^11–13^ and identified 43 rare allele carriers among 10,723 patients and 34 among 10,137 controls. To determine if the patient group includes loss-of-function alleles, we began by assaying the function of nine rare missense alleles found in patients but not in controls. These variants were also chosen to prioritize those predicted to disrupt function *in silico*. To determine the activity of LDHB variants in the most relevant context possible, we generated an *LDHB^-/-^* human iPSC line, differentiated them into motor neurons, transfected the neurons with different LDHB lentiviral expression constructs, and assayed the conversion of lactate to pyruvate catalyzed by the lysed neurons. A known LDHB loss-of-function allele, R172H^14^, was also included as a positive control. Untransfected *LDHB^-/-^* MNs have detectable but significantly lower LDH activity (48%, p=0.02) compared to isogenic control iPSC-derived MNs (‘WT’ in **Fig. 3**), demonstrating the contribution of both LDHA and LDHB to lactate metabolism in these neurons (**Fig. 3B**). Overexpression of the functional reference allele of LDHB (simply ‘LDHB’ in **Fig. 3**) in the *LDHB* KO MNs lead to markedly increased conversion of lactate to pyruvate in the assay, allowing us to compare the activities of the variant constructs. Among the initial batch of rare alleles, five showed significantly reduced activity relative to the reference allele that suggested near total loss of enzymatic function (**Fig. 3B**). One of these constructs (K23T) and the positive control R172H also reduced measured activity below that of untransfected control cells, suggesting dominant-negative effects. Thus, we identified strong loss-of-function alleles that occur among ALS cases but not controls. However, due to their rarity, we calculated that there were not enough *LDHB* missense alleles predicted to disrupt function in these data sets to allow us to generate statistical evidence that loss-of-function alleles are enriched in ALS patients. Nonetheless, our results clearly demonstrate that hemizygous congenital LDHB deficiency occurs among ALS patients, which encouraged us to employ ALS mouse models to test the hypothesis that LDHB deficiency can contribute to ALS-relevant pathology.

**Figure 3:**
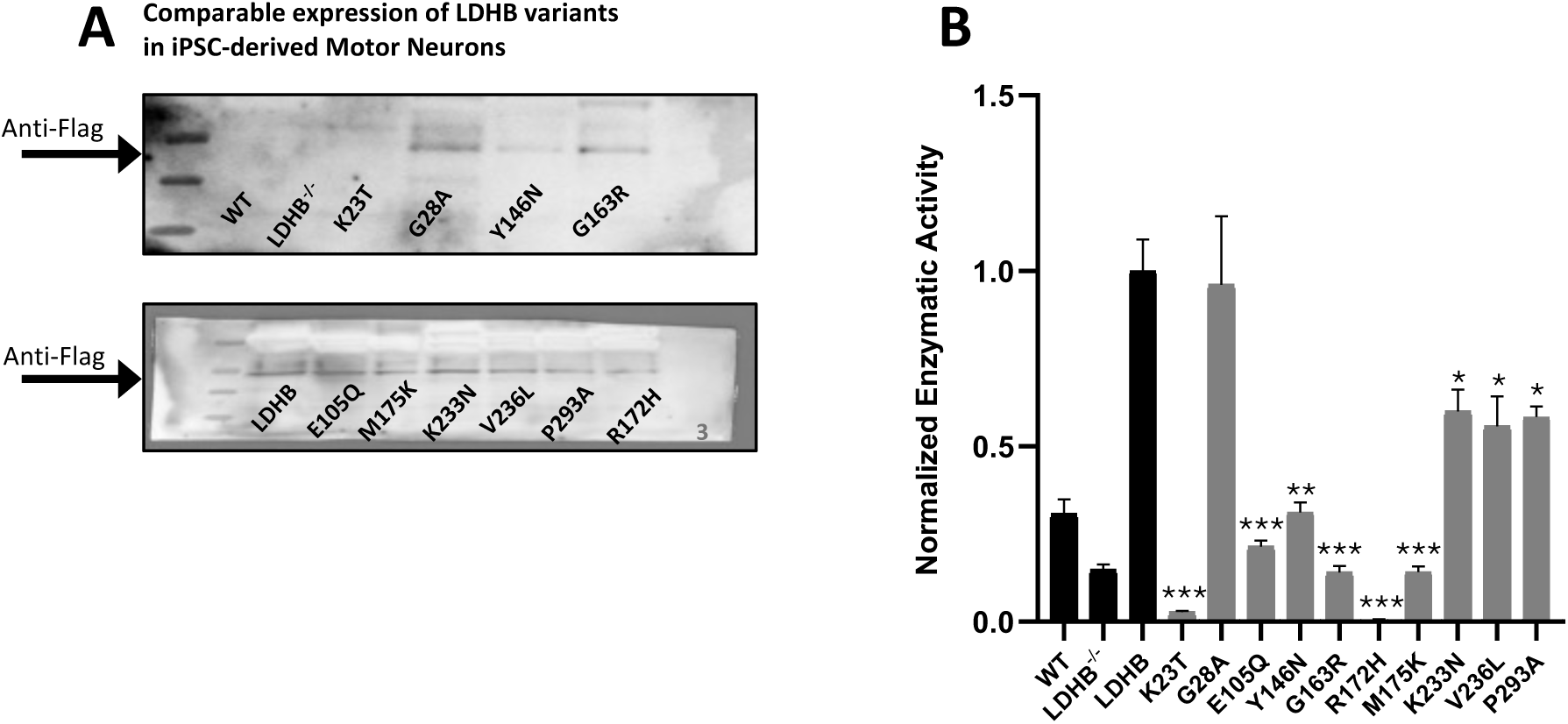
Rare *LDHB* missense alleles from ALS patients show reduced enzymatic activity in human motor neurons. Expression and enzymatic activity of the reference allele of LDHB, 9 LDHB missense alleles found in ALS patients, and R172H, a known LDHB loss-of-function allele. Western blots of motor neuron lysates probed to detect Flag-tagged LDHB protein, including untransfected controls (WT and LDHB^-/-^) **(A).** Lactase dehydrogenase activities of untransfected control (WT) and LDHB^-/-^ neurons and LDHB^-/-^ cells transfected with LDHB constructs. Enzymatic activities were calculated from the slopes of absorbance measurements normalized to the reference allele (LDHB) **(B)**. n=3 biological replicates for each construct. Statistical significance is shown for comparisons to the reference allele (LDHB) using unpaired Student’s t-test; ***p<0.001, **p<0.01, *p<0.05.

### LDHB loss in motor neurons synergizes with mildly pathogenic TDP43 to produce early motor behavior deficits

Motor neurons utilize LDHB to convert lactate to pyruvate, as demonstrated by the two-fold difference in lactate dehydrogenase activity between *LDHB^-/-^* and isogenic control MNs (**Fig. 3B**). Additionally, loss of LDHB in MNs results in mild NMJ defects as the animals age (**Fig. 2D**). Hence, *LDHB* deletion likely alters MN cellular metabolism. Reduced lactate dehydrogenase activity and lactate shuttling are associated with both normal aging and with neurodegenerative diseases, including ALS^4–6^. Therefore, we reasoned that targeted *LDHB* deletion from MNs might serve as a useful model of peripheral neurodegeneration risk due to age-associated dysregulation of lactate/pyruvate metabolism. We first explored this hypothesis by combining our MN-specific *LDHB* KO model with the TDP43^Q331K^ knock-in model^15^. TDP43 dysfunction is a hallmark of ALS, evident in >97% of cases. The TDP43^Q331K^ knock-in model carries a missense mutation in the endogenous mouse *TARDBP* locus encoding TDP43. Unlike other TDP43 ALS models that rely on overexpression of pathogenic human TDP43, this model exhibits very mild behavioral phenotypes and little evidence of pathology in the spinal cord or peripheral nerves such as TDP43 aggregation/cytoplasmic mislocalization, cell death, or axon degeneration. This allele previously proved useful to demonstrate a genetic interaction between TDP43 dysfunction and another bona fide ALS pathomechanism, STMN2 deficiency^16^. We therefore felt this model presented an appropriate sensitized background to test the potential contribution of dysregulated lactate metabolism to ALS risk.

Neither LDHB MNKO nor TDP43^Q331K/+^ mice display significant motor behavior defects when assessed by the inverted screen test up to six months of age (**Fig. 4A & S1E**), yet LDHB MNKO;TDP43^Q331K/+^ are already significantly impaired by three months (**Fig. 4A**), demonstrating notable synergy between the two sources of genetic risk. These behavioral deficits were accompanied by significant defects in NMJ innervation present in neither genetic model alone at this age (**Fig. 4E-I**). However, we did not detect significant defects in CMAP or nerve conduction velocity (NCV) **(Fig. 4B-D**). We chose to focus on the *distal* motor axon-dominated tibial nerve at this early stage because the denervation of NMJs absent axon degeneration in the LDHB KO model indicated a dying back phenotype. However, we did not observe degeneration of axons in the tibial nerves of LDHB MNKO;TDP43^Q331K/+^ mice (**Fig. 4L-P & S1A-B),** nor in their femoral nerves, a motor-enriched nerve (**Fig. S1C-D**). We also observed no evidence of TDP43 aggregation/mislocalization in the spinal cords of the combined model at six months **(Fig. 4J-K)** i.e. the synergistic phenotype cannot be explained by LDHB deficiency inducing visible TDP-43 pathology. Hence, the locus of pathology in the LDHB MNKO; TDP43^Q331K/+^ mice is likely the NMJ indicating a dying-back phenotype that has not yet progressed to frank axon degeneration, even in very distal nerves i.e. the tibial nerves.

**Figure 4:**
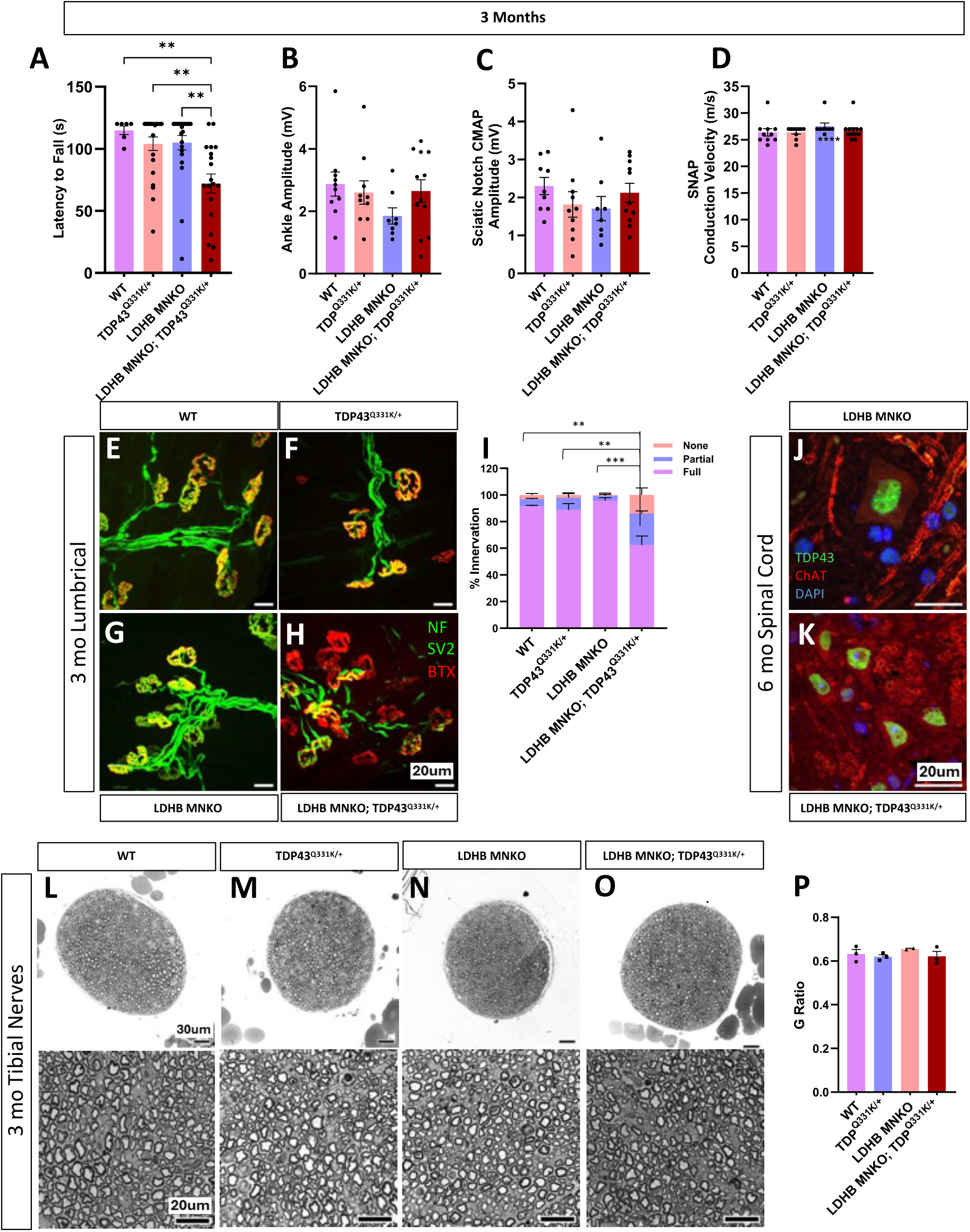
LDHB motor neuron KO synergizes with TDP43^Q331K^ to produce early motor deficits and NMJ defects: Motor function (latency to fall from an inverted screen) measured in 3-month-old WT, TDP43^Q331K/+^, LDHB MNKO and TDP43^Q331K/+^; LDHB MNKO mice **(A),** electrophysiological recording performed on 3-month-old animals to determine the difference in CMAP at the ankle and sciatic notch (**B** & **C**) and Sensory Nerve Action Potential (SNAP) at the tail (n= 10-12) **(D).** NMJs on lumbrical muscles stained with anti-NF, anti-SV2 and BTX to compare innervation of 3-month-old mice of different genotypes (n=3; statistical comparison is between percent of fully innervated) **(E-I).** Spinal cords stained to detect TDP43 and nuclei (DAPI) indicate no mislocalization in 6-month-old LDHB MNKO; TDP43^Q331K/+^ or LDHB MNKO mice **(J** & **K).** Representative images of 40x and 63x toluidine blue-stained sections of tibial nerves from 3-month-old animals (**L-O**). Average g-ratio indicates no difference in degeneration between genotypes (n= 3) **(P).** ***p<0.001, **p<0.01.

These findings are consistent with dysregulated lactate metabolism synergizing with TDP43 dysfunction to accelerate the development of a very distal neuropathy. These findings are consistent with a model in which age- or disease-related decline in lactate/pyruvate metabolism influences the timing of ALS initiation or progression.

### LDHB loss in motor neurons synergizes with mildly pathogenic SOD1 to produce early motor behavior deficits

To test whether the synergy we observed between LDHB loss and pathogenic TDP43 is specific to this pathomechanism or is more general to ALS pathology, we chose to investigate the contribution of dysregulated lactate metabolism in another distinct ALS model. Similar to the *TDP43^Q331K^* knock-in model, the *Sod1^D83G^*model does not rely on overexpression, in contrast to the commonly investigated SOD1-G93 model, which massively overexpresses pathogenic human SOD1^17^. Instead, *Sod1^D83G^* is a spontaneous point mutation identical to a human familial ALS-associated *SOD1* variant with notably variable penetrance^18,19^. Progressive motor defects occur reliably in the *Sod1^D83G^* mouse line, but only in homozygous males relatively late in life^18^. We therefore saw this model as an ideal alternative sensitized background to test the generalizability of the disease risk associated with dysregulated lactate metabolism. Similar to the TDP43 model, combining LDHB MNKO with *Sod1^D83G/D83G^*also led to significantly earlier and more severe motor deficits (**Fig. 5**), demonstrating that dysregulated lactate metabolism is a significant motor neurodegeneration risk synergizing with ALS risk factors acting via disparate mechanisms.

**Figure 5:**
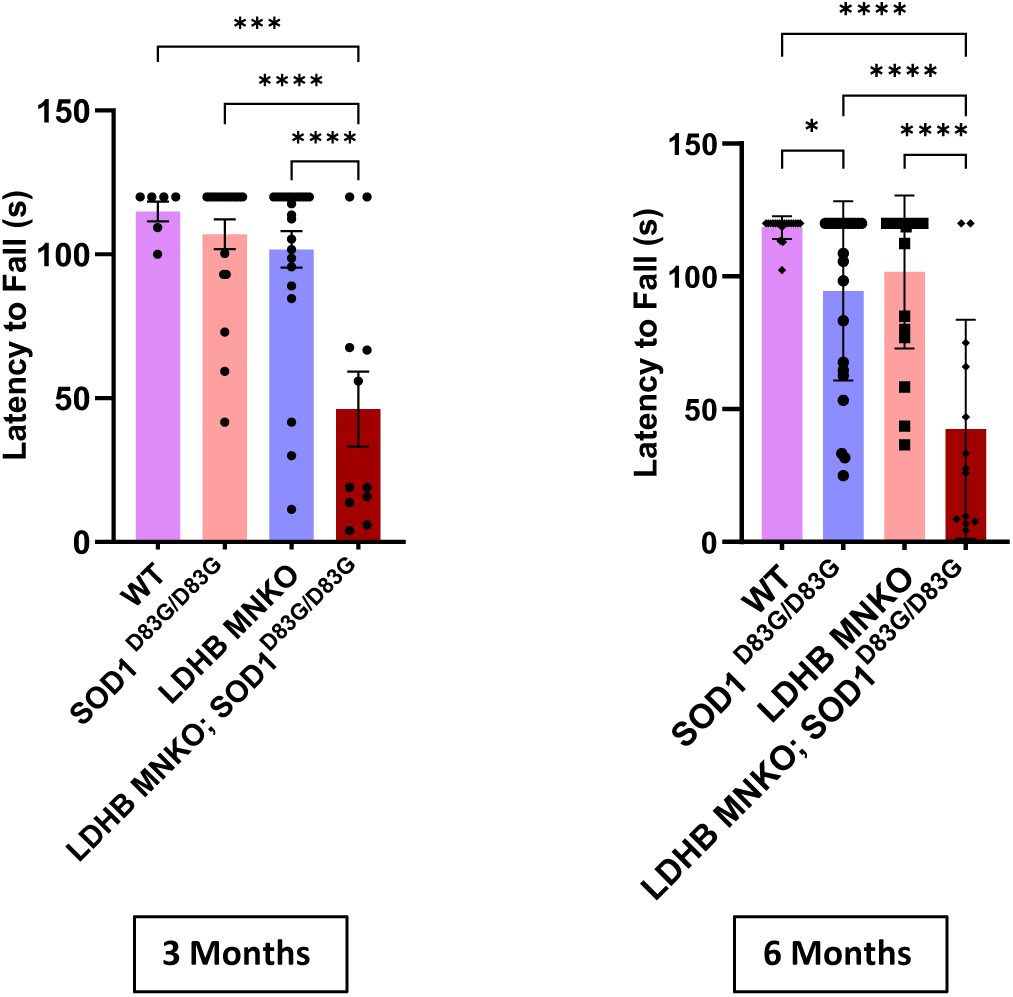
LDHB MNKO synergizes with SOD1^D83G^ to produce defects in motor behavior: Motor function (inverted screen) of 3 and 6-month-old WT, LDHB MNKO, SOD1^D83G/D83G^ and LDHB MNKO; SOD1^D83G/D83G^ mice (n= 15-20). ****p<10^-4^, ***p<0.001.

## Discussion

While significant advances have been made in understanding ALS etiology by studying highly penetrant pathogenic variants in genes like SOD1, TARDBP and C9ORF72, most sporadic ALS cases are understood to arise from the intersection of lesser risk factors, both congenital and environmental, including genetic modifiers that influence disease onset, progression, and penetrance^20,21^. Importantly, the best predictor of ALS is advanced age^22^, which is associated with the decline of numerous cellular and intercellular processes including lactate/pyruvate metabolism^4,23^. Lactate production in the CNS declines with age, and deteriorated lactate shuttling from glia to neurons is implicated in aging-related neurodegenerative diseases, including Alzheimer’s and ALS^4–6^. Evidence for this includes metabolomic studies of sporadic ALS patient serum that documented disrupted lactic acid metabolism^24^. By ablating lactate dehydrogenase activity in Schwann cells, we previously demonstrated that a similar lactate shuttling system is also necessary to maintain motor axons but not sensory axons in the PNS, suggesting a connection between compromised lactate metabolism and motor-selective peripheral neuropathy risk^8^. Here we have shown that, while motor neurons employ both LDH subunits, LDHB is not necessary to maintain motor axons, and that early motor defects displayed by whole-body LDHB knockout mice can be attributed to LDHB deficiency in Schwann cells rather than neurons. Combining murine models to assess genetic interactions contributing to ALS risk has previously proven fruitful^16,25^. Therefore, to interrogate the risk for peripheral neurodegeneration due to dysregulated lactate metabolism separate from other age-related changes, we utilized a genetic model of LDH deficiency by knocking out only LDHB (not LDHA) in motor neurons. Tellingly, although this mild disruption has little effect on motor function by itself, in combination with other weak ALS genetic risk factors, striking declines in motor function are observed. Thus, altered lactate utilization by motor neurons can significantly exacerbate pathogenesis, suggesting that lactate metabolism function could significantly impact disease susceptibility and progression.

In a prior study, we combined ALS case–control exomes with functional assays to demonstrate that a subset of rare variants in the *SARM1* gene confer constitutive NADase activity and are greater than 5-fold enriched in ALS patients, supporting SARM1 gain-of-function as a genetic risk mechanism for ALS^10^. By contrast, in this study, driven by results from mouse models, we analyzed LDHB assuming the more typical loss-of-function axis of risk. Using an LDHB^-/-^ iPSC motor neuron background, we functionally tested rare LDHB variants observed in ALS datasets and identified multiple severe loss-of-function alleles (sometimes with apparent dominant-negative behavior) present among ALS patients but absent from controls. However, because such alleles are very rare, our current sample does not provide statistical evidence for case enrichment. Nevertheless, taken together with our mouse genetics results, the human data are consistent with LDHB haploinsufficiency acting as a disease modifier that sensitizes motor units to otherwise subthreshold ALS risks if not a stand-alone causal factor. Limitations of our analysis include the rarity and ascertainment of LDHB variants in current exome cohorts and the constraints of our overexpression-based assay. Larger meta-analyses and systematic burden tests will be needed to determine whether LDHB loss-of-function is truly enriched in ALS at the population level.

Axons vary in their metabolic requirements in parallel with their diverse roles and morphologies. Spinal motor axons are exceptionally long and elaborate, extending up to a meter in humans and innervating muscles via numerous NMJs. Synapses expend more than a third of a neuron’s ATP production, which is regenerated locally via mitochondrial oxidative phosphorylation^26,27^. Thus, the demands of propagating action potentials over long distances and fueling large numbers of synapses confer unique energy requirements on spinal MNs that likely contribute to their special vulnerability in ALS^28^. Indeed, the MNs that innervate fast-twitch muscle and have the highest peak ATP requirements are also those affected the earliest and most severely in ALS; MNs that innervate slow-twitch muscle are affected later, and oculomotor neurons, which are exceptionally resistant to ALS, innervate relatively few NMJ endplates compared to spinal MNs ^29^. As such, the motor-selectivity of neuropathy induced by LDH knockout fits with a model of vulnerability related to high metabolic stress within the neuron itself.

Mitochondrial ATP production is impaired in ALS^30^, and many mitochondrial genes are associated with ALS and other peripheral neuropathies^31^. Pathogenic variants of SOD1 and TDP-43 both impair mitochondrial function and motor neurons derived from ALS patient iPS cells display elevated LDH leakage along with reduced mitochondrial respiration and ATP production^32^. We propose the following model as a potential explanation for our findings of synergy. Altered cellular pyruvate supply in the background of a compromised TCA cycle might overwhelm compensatory mechanisms and cross a tipping-point to produce metabolic deficiency worse than either insult alone. Such synergy could accelerate pathology due to oxidative stress as reductive oxygen species (ROS) byproducts are generated by incomplete substrate oxidation in impaired and inefficient mitochondria. Thus, even when other cellular defects occur, boosting lactate metabolism/shuttling to cushion against the consequences of poor mitochondrial function could protect axons and forestall pathology ^33,34^. In short, our results suggest that impaired lactate metabolism can accelerate pathology in ALS models, a compelling rationale to purse lactate restoration as a therapeutic strategy for peripheral neuropathies.

## Methods

### Animals

All animal experiments were performed with the approval of the Institutional Animal Care and Use Committee of Washington University, St. Louis, MO. Floxed Ldha (*Ldha^F/F^*) mice were originally generated by breeding *Ldha*^tm1a(EUCOMM)Wtsi^ mice (EMMA, EM:05082, Jax Stock No: 030112)^35^ to mice that express FLP recombinase ACTB:FLPe B6;SJL (Jax: 003800)^36^. Floxed Ldhb (*Ldhb^F/F^*) mice were similarly generated from Ldhb^tm1a(KOMP)Wtsi^ mice (EMMA, EM:08936)^37^. To generate motor neuron and Schwann cell specific LDHB knockout mice, *Ldhb^F/F^* mice were bred to either ChAT-Cre^+^ or MPZ-Cre^+^ mice^38^ (Jax: 006410)^39^ respectively. Whole-body Ldha and Ldhb KO lines were generated by mating *Ldha^F/F^* or *Ldhb^F/F^* mice to mice expressing Cre driven by the actin promoter. TDP-43^Q331K^ (Jax:031345) and Sod1^D83G^ (Jax:020440) mice were obtained from Jackson Laboratories. Littermate controls consisted of Cre negative mice in all experiments. Male and female mice were used in approximately equal numbers in all experiments except where otherwise specified.

### Inverted Screen test

The inverted screen test was performed as described previously^16^. Mice were set on a wire mesh which was then inverted for a maximum of 2 minutes. Each mouse was tested thrice at an interval of at least 5 minutes. The latency to fall was calculated based on the average of three measurements.

### Nerve electrophysiology

Compound muscle action potential (CMAP) and sensory nerve action potential (SNAP) were acquired using a Viking Quest electromyography device (Nicolet) as we previously published^42^. Mice were anesthetized with isoflurane, with body temperature maintained at 37 °C with a thermostatic heating pad and rectal probe, and CMAP for ankle and sciatic notch was recorded by inserting the recording electrodes in the plantar surface of the footpad (intrinsic foot muscles) and the stimulating electrode in the ankle (distal) or sciatic notch (proximal), respectively. The ground and reference electrodes were inserted subcutaneously in the tail base. Both hindlimbs were recorded using supramaximal (≥20% above the plateau CMAP amplitude) stimulation for CMAPs and 3-5 traces were averaged for each site. Supramaximal stimulation was also used for SNAP measurements. Electrodes were inserted subcutaneously into the tail. The recording electrode was placed at the base of the tail, followed by the ground, the stimulating, and the reference electrode with a fixed 30 mm distance between the recording and stimulating electrode.

### Nerve structure analysis using light microscopy

As described in our previous work ^16^ nerves were fixed in 3% Glutaraldehyde overnight at 4°C, washed, and stained in 1% Osmium Tetroxide overnight at 4°C. These nerves were washed and dehydrated in a serial gradient of ethanol. After dehydration, nerves were incubated in 50% propylene oxide/50% ethanol, then 100% propylene oxide. Subsequently, nerves were incubated in Araldite resin solution/propylene oxide solutions overnight, and then embedded in 100% Araldite resin solution (Araldite: DDSA: DMP30; 12:9:1; Electron Microscopy Sciences) and finally baked overnight at 60°C. For the light microscope analysis, sections of 400–600 nm were cut using Leica EM UC7 Ultramicrotome and placed onto prewarmed slides and stained for 1 minute with 1% toluidine blue solution (1% toluidine blue, 2% borax), vigorously washed with water, acetone, and then xylene. They were mounted with Cytoseal XYL and imaged on a compound brightfield microscope. Images taken with either a 100x or 63x oil lens were used for quantification. g-ratio was measured by dividing the inner myelin diameter by outer diameter, quantifying 40 myelinated axons per mouse and averaging across replicates. Degenerated axons were excluded from g-ratio calculations.

### NMJ staining and analysis

Mice were transcardially perfused with 4% Paraformaldehyde (PFA) followed by overnight fixation of the feet in 4% PFA. Following three rinses with Phosphate buffered saline (PBS), lumbrical muscles were dissected for staining with anti-SV2 (Developmental Studies Hybridoma Bank AB2315387, 1:200), anti-2H3 (Developmental Studies Hybridoma Bank AB2314897, 1:100) and α-bungarotoxin (BTX) (Biotium 00006, 1:500) as described previously^8^. NMJ images were acquired at 63x and 40x magnification on a confocal microscope using the z-stack setting. Image analysis was performed by projecting these images at maximum intensity, followed by observing the colocalization of the pre-synapse with the post-synapse. The individual NMJs were categorized as fully, partially, or not innervated by an experimenter blinded to genotype. The proportion of innervation was translated into percentages and compared across genotypes by 2-way ANOVA followed by multiple comparison testing.

### Spinal Cord staining

Mice were transcardially perfused with 4% PFA, and their spinal columns were dissected out and fixed further for 48 hours in 4% PFA. They were then immersed in 30% sucrose and stored at 4°C until dissection. Spinal cords were dissected, transferred into 70% ethanol, and sliced into six pieces for embedding in paraffin using a tissue processor. The embedded paraffin blocks were sliced using a microtome, and 5 µm sections were placed on prewarmed slides. Sections were stained using antigen retrieval with citrate buffer and pressure-cooked for 2 minutes. The slides were cooled at room temperature for 30 minutes and gently washed with water before blocking in 4% BSA + 0.3% Triton-X100 in 1X PBS for 30 minutes. They were incubated with Rabbit anti-ChAT (Millipore A. B143) antibody at 4°C overnight. The next day, slides were washed with 0.03% Triton X-100 in PBS and incubated with C-terminal conjugated anti-TDP43 (Proteintech CoraLite CL488- 67345) along with Goat anti-rabbit IgG (H+L) cross-adsorbed secondary antibody, Alexa Flour 568 (Invitrogen A11011). The stained slides were mounted with Vectashield containing DAPI and imaged on a confocal microscope at 63x magnification using the Z-Stack setting. Images were observed for TDP43 mislocalization by projecting for maximum intensity.

### iPSC Differentiation into Motor Neurons

LDHB knockout iPSCs were generated by the Genome Engineering and iPSC Center (GEiC) at Washington University in St. Louis. Human iPSCs were differentiated into spinal motor neurons using a previously established protocol^40^ with minor adjustments. Neural induction was initiated with small-molecule inhibitors, followed by patterning with retinoic acid and purmorphamine to generate OLIG2⁺ motor neuron progenitors (MNPs). MNPs were expanded and matured into functional CHAT⁺ motor neurons with valproic acid, Compound E, and neurotrophic factors, including IGF-1, BDNF, and CNTF.

### LDHB Variant Activity Assay from Cell Lysates

Synthetic DNA fragments encoding human LDHB were cloned into the lentivirus vector FCIV using InFusion (Clontech) and lentiviral particles containing LDHB variants were generated as previously described^41^. Variants were selected from anonymized publicly available human genetic data not associated with demographic or clinical information beyond ALS case status. All assayed polymorphisms were reported in one of three databases, last accessed July 2024: Project MinE (http://databrowser.projectmine.com)^11^, the University of Massachusetts Medical School Sporadic ALS Variant Server (http://als.umassmed.edu/ index.php#SALSbrowser), or the ALS Knowledge Portal (http://alskp.org)^12^. LDHB knockout (KO) motor neuron precursors derived from human induced pluripotent stem cells (iPSCs) were transduced with lentivirus expressing LDHB variants (n= 3 biological replicates per construct). Following differentiation into mature motor neurons, equal amounts of protein from each cell lysate were added to each assay and LDH activity was measured using the Lactate Dehydrogenase Activity Assay Kit (Millipore Sigma, MAK066) detecting absorbance at 450 nm every 5 minutes for one hour. Enzymatic activities were calculated from the slope of the reaction curves with all activities represented relative to LDHB KO cells overexpressing the LDHB reference allele construct normalized to 1.

### Statistical Analysis

Sample numbers (n) of ≥3 were used in all experiments, with numbers specified throughout. Statistics were performed using GraphPad Prism. All data is reported as mean ± SEM. All group comparisons were performed using one-way ANOVA followed by multiple comparison tests, except for the percentage of innervation, which was analyzed using two-way ANOVA followed by multiple comparison tests.

## Acknowledgments

We would like to thank members of the DiAntonio and Milbrandt labs for their thoughtful feedback on this work. We would like to thank Cassidy Menendez and Liya Yuan for their technical support.

---

**Supplementary Figure 1.**
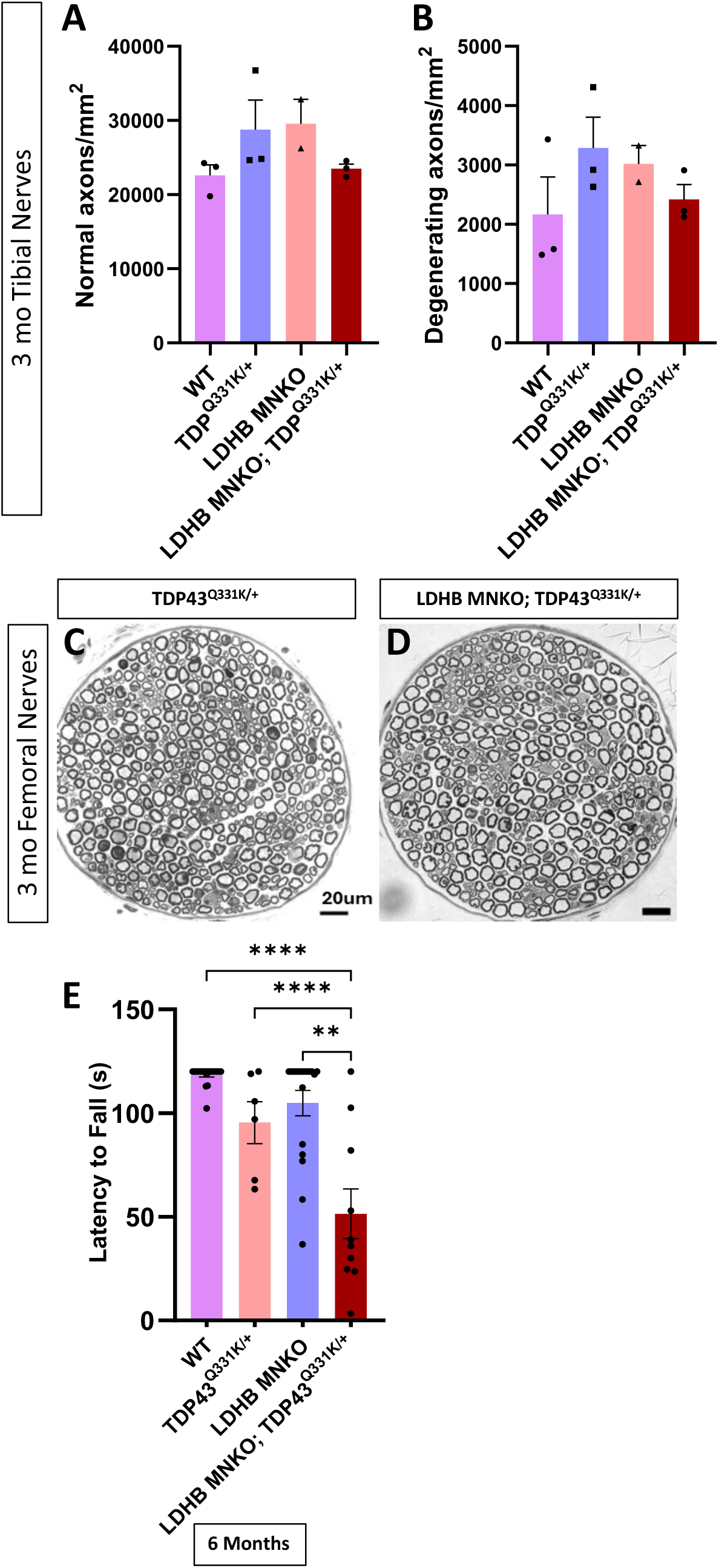
Density of normal and degenerating axons in tibial nerves of WT, TDP43^Q331K/+^, LDHB MNKO, and LDHB MNKO; TDP43^Q331K/+^ mice **(A** & **B).** Representative images of 63x toluidine blue-stained sections of Femoral nerves from 3-month-old TDP43^Q331K/+^ and LDHB MNKO; TDP43^Q331K/+^ mice **(C** & **D).** Latency to fall from an inverted screen in 6-month-old animals **(E).** ***p<0.001, **p<0.01.

